# Hypoxic conditions promote *Candida glabrata* colonization in the intestinal tract and *EPA6* plays a significant role in hypoxic adhesion to intestinal cells

**DOI:** 10.1101/2022.07.01.498396

**Authors:** Takayuki Shinohara, Masahiro Abe, Sota Sadamoto, Minoru Nagi, Harutaka Katano, Hiroji Chibana, Yoshitsugu Miyazaki

## Abstract

*Candida glabrata*, a fungal pathogen colonizing mucocutaneous membranes and indwelling medical devices, is associated with invasive infections, especially in immunocompromised individuals. Candidiasis could be of endogenous and exogenous origins. Endogenous infections are considered to derive from the invasion of *Candida* species colonizing the digestive mucosa. Investigations of the gut-to-bloodstream translocation mechanisms of *Candida* species remain limited, although environmental oxygen levels have been recently suggested to alter the human fungal pathogen phenotypes. Moreover, human fungal pathogens, including *Candida*, colonizing or invading less oxygenated tissues encounter altered oxygen circumstances. Therefore, phenotype investigation under hypoxic conditions could provide valuable novel insights into the host-pathogen interaction mechanisms. This study aimed to elucidate the adhesion capabilities and mechanisms of *C. glabrata* depending on various oxygen levels. We performed *C. glabrata* adhesion assays to Caco-2 cells under aerobic, microaerobic (5 vol% oxygen), and anaerobic conditions, conducted RNA-seq to identify candidate genes functioning on hypoxic adhesion. We then generated deletants of these genes and evaluated both their adhesion to Caco-2 cells under anaerobic conditions and their colonization ability in the hypoxic intestinal tract in a mouse model. We observed significant differences in Caco-2 cell adhesion in response to different oxygen levels. Under hypoxic conditions, the *C. glabrata* adhesion capability increased and the expression levels of seven adhesion-related genes were up-regulated. Among these mutants, the adhesion capability of *epa6*Δ decreased the most. The *epa6*Δ mutant exhibited significantly lower intestinal colonization in mice than the wild-type. To the best of our knowledge, this study first describes the hypoxic adjustment of *C. glabrata* to intestinal cell adhesion, in which *EPA6* plays the most significant role. If Epa6p function could be inhibited, it may contribute to reducing endogenous infection. Phenotype investigation under hypoxic conditions could provide valuable novel insights into the host-pathogen interaction mechanisms.

**Author Summary:** *Candida glabrata* is the second most common pathogen of *Candida* infections (i.e., candidiasis), colonizing mucocutaneous membranes, indwelling medical devices, thereby causing bloodstream- and medical device-related infections and often leading to high morbidity and mortality. Candidiasis could be of endogenous and exogenous origins. Endogenous infections are considered to derive from the invasion of *Candida* species colonizing the digestive mucosa. Investigations of the gut-to-bloodstream colonizing and translocation mechanisms of *Candida* species remain limited. Interestingly, recent studies suggest that environmental oxygen levels could alter the human fungal pathogen phenotypes. This study thus focused on the relationship between the colonization and adhesion capability of *C. glabrata* in the gastrointestinal tract depending on the environmental oxygen level to address the underlying mechanisms. Our results suggest that anaerobic conditions promote *C. glabrata* adhesion and *EPA6* plays a significant role in hypoxic adhesion, opening new perspectives in various affiliated fields and related research domains. If Epa6p function could be inhibited, it may contribute to control the colonization in the gut and following translocation. *C. glabrata* is known to be low-susceptible to azole antifungals. A novel antifungal agent type, such as one targeting these adhesive molecules, should thus be considered and further related studies would be necessary.

## Introduction

*Candida glabrata*, opportunistic fungus in the mucosal flora, can cause bloodstream- and medical device-related infections [1]. Immunosuppressed states, indwelling devices (such as catheters), parenteral nutrition, and antibiotic overuse are common contributing factors to *Candida* infections or candidiasis [2]. While *Candida albicans* remains the leading cause of such infections, *C. glabrata* is the second most common pathogen of candidiasis, accounting for 15–20% of disseminated candidiasis [3]. *Candida* infections often lead to high morbidity and mortality [4], exploring factors and molecular determinants for *Candida* pathogenicity is thus an urgent challenge.

Candidiasis could be of endogenous and exogenous origins. Endogenous infections are considered to derive from the invasion of *Candida* species colonizing the digestive mucosa [5]. Investigations of the gut-to-bloodstream colonizing and translocation mechanisms of *Candida* species remain limited. Environmental parameters including temperature, pH, serum, and CO_2_ are associated with several steps during the host invasion and optimal growth of *Candida* species [6]. Hypoxic conditions are recently considered to be important factors potentially affecting the virulence of the pathogen [6]. In the gut, the residence of various indigenous or pathogenic microorganisms, oxygen concentrations range between 0–70 mmHg, and similar concentrations occur in the liver, pancreas, and peritoneal cavity [7]. Of note, human fungal pathogens, including *Candida* species, colonizing or invading less oxygenated tissues encounter alterations in oxygen availability [8]. However, studies focusing on fungal phenotypes under hypoxic conditions are scarce.

Regarding *C. albicans*, the morphological transformation of yeast cells into elongated budding tubes enhances their ability to invade tissues and contribute to the lesional expansion, regulated by the transcription factor *EFG1* [9]. Subsequent studies showed that the regulator protein Efg1p represses filaments under hypoxic conditions [10–12], favoring commensalism in the hypoxic niches of the human host [13, 14]. Moreover, favorable oxygen conditions for biofilm formation reportedly differ between *Candida* species [15]. *Candida tropicalis* promotes hyphal formation under microaerobic (5 vol% oxygen) compared to aerobic and anaerobic conditions [15]. We thus consider that the *C. glabrata* phenotype can also change in response to oxygen concentration. Phenotype investigation under hypoxic conditions could provide insights into the host-pathogen interaction mechanisms.

In this study, related to endogenous *C. glabrata* infection, we focused on the relationship between the colonization capability of *C. glabrata* in the gastrointestinal tract and the environmental oxygen level and evaluated *C. glabrata* adhesion to intestine-derived Caco-2 cells under various oxygen concentrations. We observed that the adhesion capability increased under hypoxic conditions. Based on this finding, we conducted RNA-seq analysis to identify potent adhesion genes induced under hypoxic conditions. We generated deletants of these genes and evaluated not only their adhesion to Caco-2 cells under anaerobic conditions but also their colonization ability in the intestinal tract using a murine model.

## Results

### Anaerobic conditions increased the adhesion capability to Caco-2 cells of *C. glabrata*

We first investigated whether oxygen concentration, i.e., aerobic (21 vol% oxygen), microaerobic (5 vol% oxygen), and anaerobic (0 vol% oxygen) conditions influenced the adhesion capability of *C. glabrata*. Adhesion assays on Caco-2 cells were performed for the CBS138 reference strain and four clinical strains isolated from bloodstream infections. We observed significant differences in the adhesion capabilities of each strain leading to an augmented response to decreasing oxygen concentrations (Fig 1).

**Fig 1.**
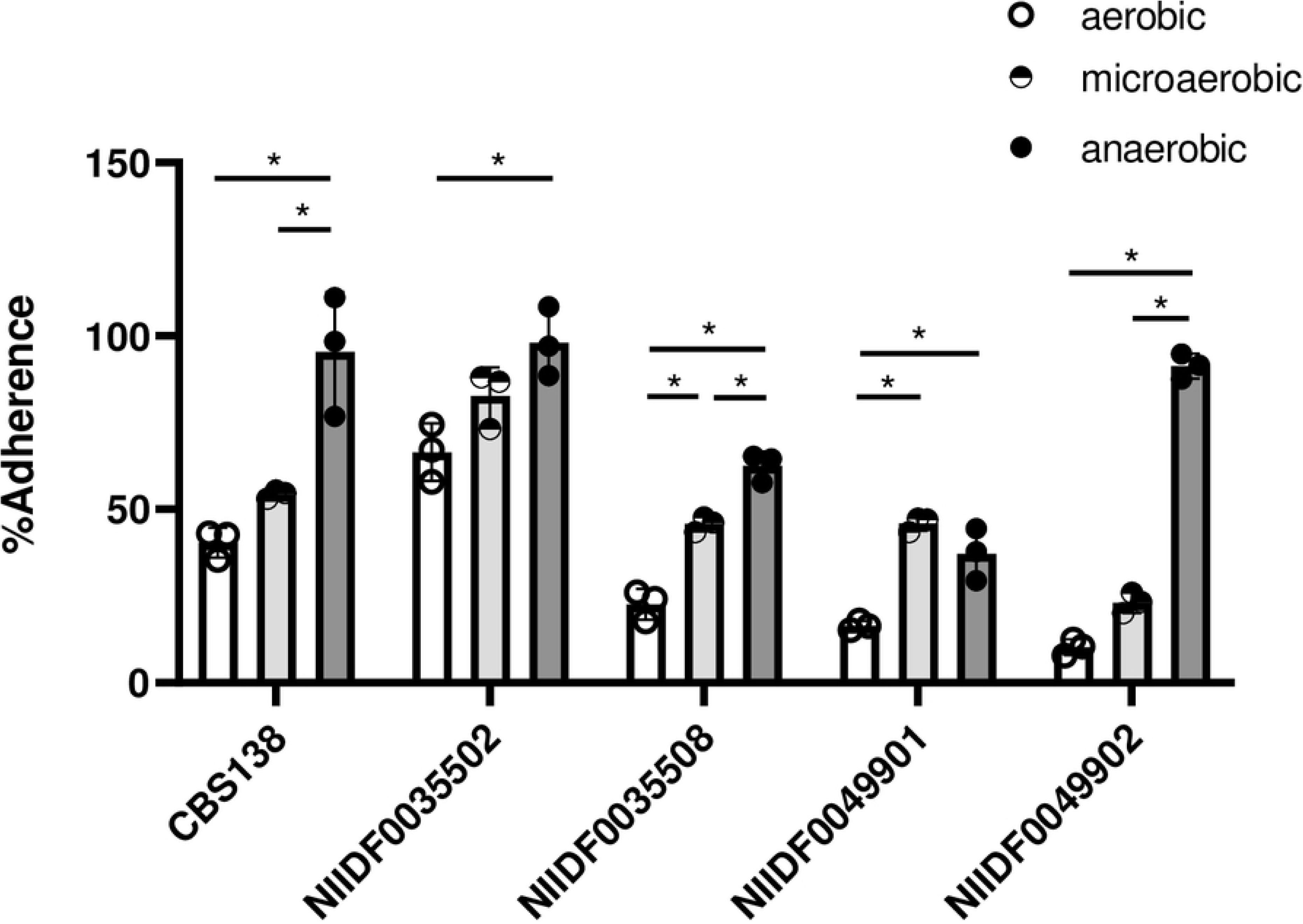
The adhesion capability of *C. glabrata* strains to Caco-2 cells increased under hypoxic conditions. NIIDF0035502, NIIDF0035508, NIIDF0049901, and NIIDF0049902 were clinical strains isolated from bloodstream infections. All results are expressed as the mean ± standard error of the mean from three independent experiments. * *p* < 0.05.

### RNA-seq uncovered seven *C. glabrata* adhesins involved in anaerobic adhesion

The increased adhesion capability of *C. glabrata* under anaerobic conditions suggested that certain adhesion gene expressions could be up-regulated during anaerobic adhesion. Therefore, we applied RNA-seq to explore adhesion genes up-regulated in anaerobic adhesion. We compared the TPM values of *C. glabrata* extracted at the time of inoculum and two hours after Caco-2 cell adhesion under anaerobic and aerobic conditions, respectively (S2 Table presents complete datasets). Our results identified 711 genes (Group A) more than two-fold up-regulated upon two hours of anaerobic adhesion (Fig 2A, red dots), and 1,339 genes (Group B) changed less than two-fold upon two hours of aerobic adhesion (Fig 2B, blue dots). We discovered 355 genes common to Group A and B (Fig 2C), including seven adhesion genes: *EPA2, EPA6, AWP1, AWP2, PWP3, CAGL0J00253g*, and *CAGL0J02552g*. S3 Table shows the TPM value fold changes of adhesion genes upon two hours under aerobic and anaerobic adhesion. A further analysis was performed to identify the genes exerting the greatest influence on anaerobic adhesion. Fig 2D shows the ratios of TPM value fold changes upon two hours of anaerobic adhesion divided by those upon two hours of aerobic adhesion, listed in descending order. These results indicated that *EPA6* was more up-regulated under anaerobic compared to aerobic adhesion.

**Fig 2.**
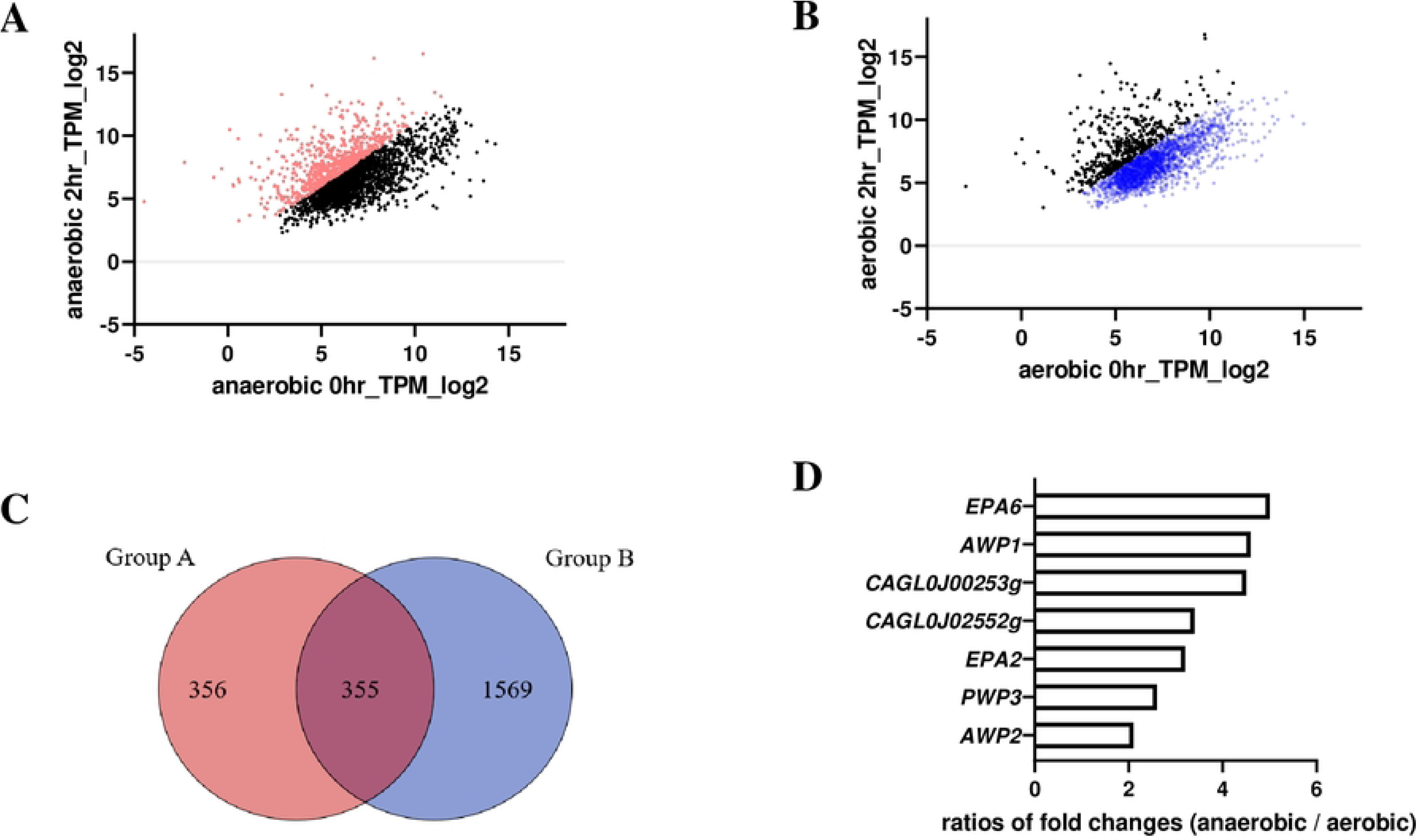
Gene expression of *C. glabrata* upon two-hour adhesion to Caco-2 cells under aerobic and anaerobic conditions. (A and B) Scatter plots of gene TPM values (in log2) measured by RNA-seq in anaerobic and aerobic adhesion to Caco-2 cells. More than two-fold up-regulated under anerobic upon two hours and less than two-fold changed under aerobic upon two hours are represented by red and blue dots, respectively. (C) Venn diagram. Group A consisted of 711 genes that were up-regulated more than two-fold in anaerobic adhesion. Group B consisted of 1924 genes that were changed less than two-fold in aerobic adhesion. Group A and B contained 355 common genes common. (D) Ratios of anaerobic-to-aerobic fold change in seven adhesion genes.

### *EPA6* governed *C. glabrata* adhesion to Caco-2 cells under anaerobic conditions

We generated *C. glabrata* mutants for seven adhesion genes (*EPA2, EPA6, AWP1, AWP2, PWP3, CAGL0J00253g*, and *CAGL0J02552g*) involved in anaerobic adhesion and performed adhesion assays on Caco-2 cells under anaerobic conditions. Each target gene was disrupted by a DNA replacement cassette including the *CgHIS3* gene through homologous recombination, otherwise, *CAG0J02552g* was suppressed using a Tet-OFF promoter system as described previously [16]. Of the seven mutants, *epa6*Δ showed the most reduced adhesion capability (Fig 3A) with a statistically significant difference compared to the parent strain (Fig 3B). In the reintegrated strain (*epa6Δ/EPA6*), adhesion was significantly recovered (Fig 3B).

**Fig 3.**
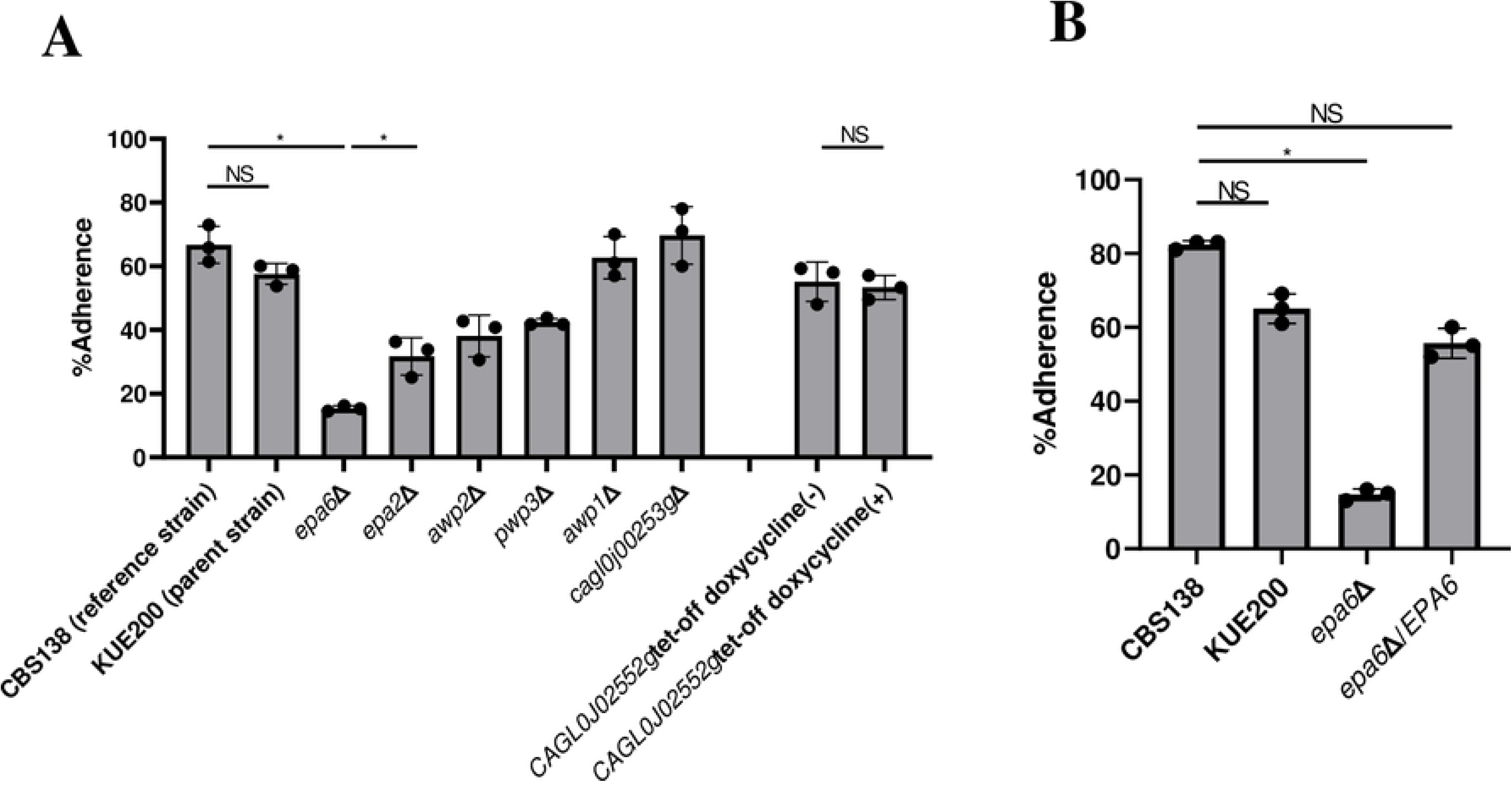
The *epa6*Δ mutant showed the most reduced adhesive capability to Caco-2 cells under anaerobic conditions. (A) Adhesion assays were performed on CBS138 (reference strain), KUE200 (parent strain), and seven mutants. The *CAG0J02552g* Tet-OFF mutant was evaluated with or without 10 μg/ml of doxycycline in the medium from culture to adhesion assays. All results are expressed as the mean ± standard error of the mean from three independent experiments. (B) Adhesion assays and significant tests were performed on CBS138, KUE200, *epa6*Δ, and *epa*6Δ/*EPA6*. All results are expressed as the mean ± standard error of the mean from three independent experiments. * *p* < 0.05, NS, no significant difference.

### *EPA6* transcript level was up-regulated early under anaerobic adhesion to Caco-2 cells

We investigated how the *EPA6* expression level changed over time using RT-qPCR in the wild-type (CBS138), parent (KUE200), and *EPA6* reintegrated (*epa6*Δ/*EPA6*) strains at hour zero (start of adhesion to Caco-2 cells), followed by two and eight hours of adhesion. We found that the *EPA6* mRNA accumulated approximately 40-fold higher at two hours of anaerobic adhesion to Caco-2 cells than that at hour zero (Fig 4). However, the *EPA6* transcript level did not change significantly in aerobic adhesion.

**Fig 4.**
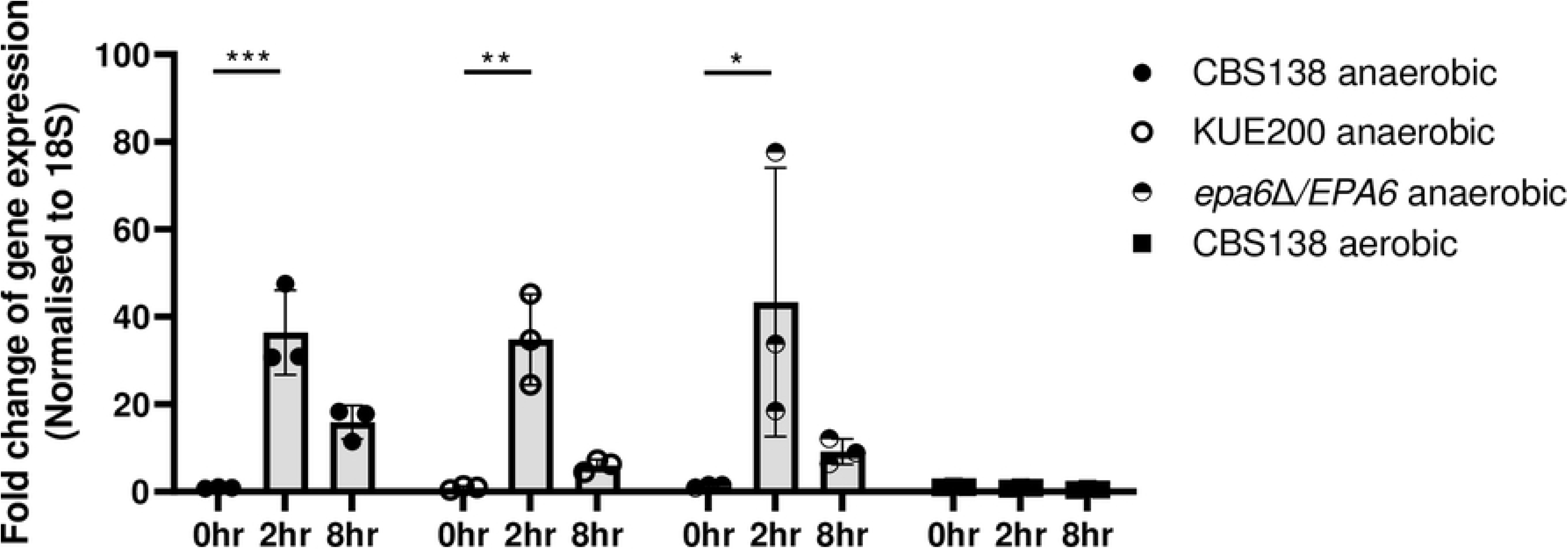
The *EPA6* transcript level changed drastically under anaerobic conditions. cDNA was prepared using total RNA samples from strains at 0-, 2-, and 8-hour adhesion to Caco-2 cells under anaerobic or aerobic conditions. Quantitative RT-PCR analysis was performed, and the expression level of each strain was exhibited as relative fold changes compared to wild-type (CBS138) cells at hour zero (the start of adhesion). All results are expressed as the mean ± standard error of the mean from three independent experiments. * *p* < 0.05, ** *p* < 0.01. *** *p* < 0.001.

### *C. glabrata EPA6* disruption resulted in reduced gastrointestinal colonization in mice

To test the correlation between in vivo and in vitro adhesion via Epa6p, we set up a murine model of *C. glabrata* colonization in the gut. We intragastrically inoculated each mouse with the *C. glabrata* CBS138, *EPA6* deletant (*epa6*Δ), and reintegrated (*epa6*Δ/*EPA6*) strains. After inoculation, we evaluated the gastrointestinal colonization of each group by measuring stool CFUs. The fungal burden in the gastrointestinal tract was significantly lower in mice infected with *epa6*Δ than in those infected with CBS138 and *epa6*Δ/*EPA6* 7 days after inoculation (Fig 5). In addition, based on this result, we histopathologically evaluated the guts 7 days after inoculation with CBS138, *epa6*Δ, and *epa6*Δ/*EPA6*. The histological analyses revealed that round yeasts, albeit a few per high-power field, were present in the intestine of mice infected with CBS138 and *epa6*Δ/*EPA6* (Fig 6A and C), however, they were scarcely in the case of *epa6*Δ infection (Fig 6B). These results indicate that Epa6p could be of critical importance for intestinal colonization.

**Fig 5.**
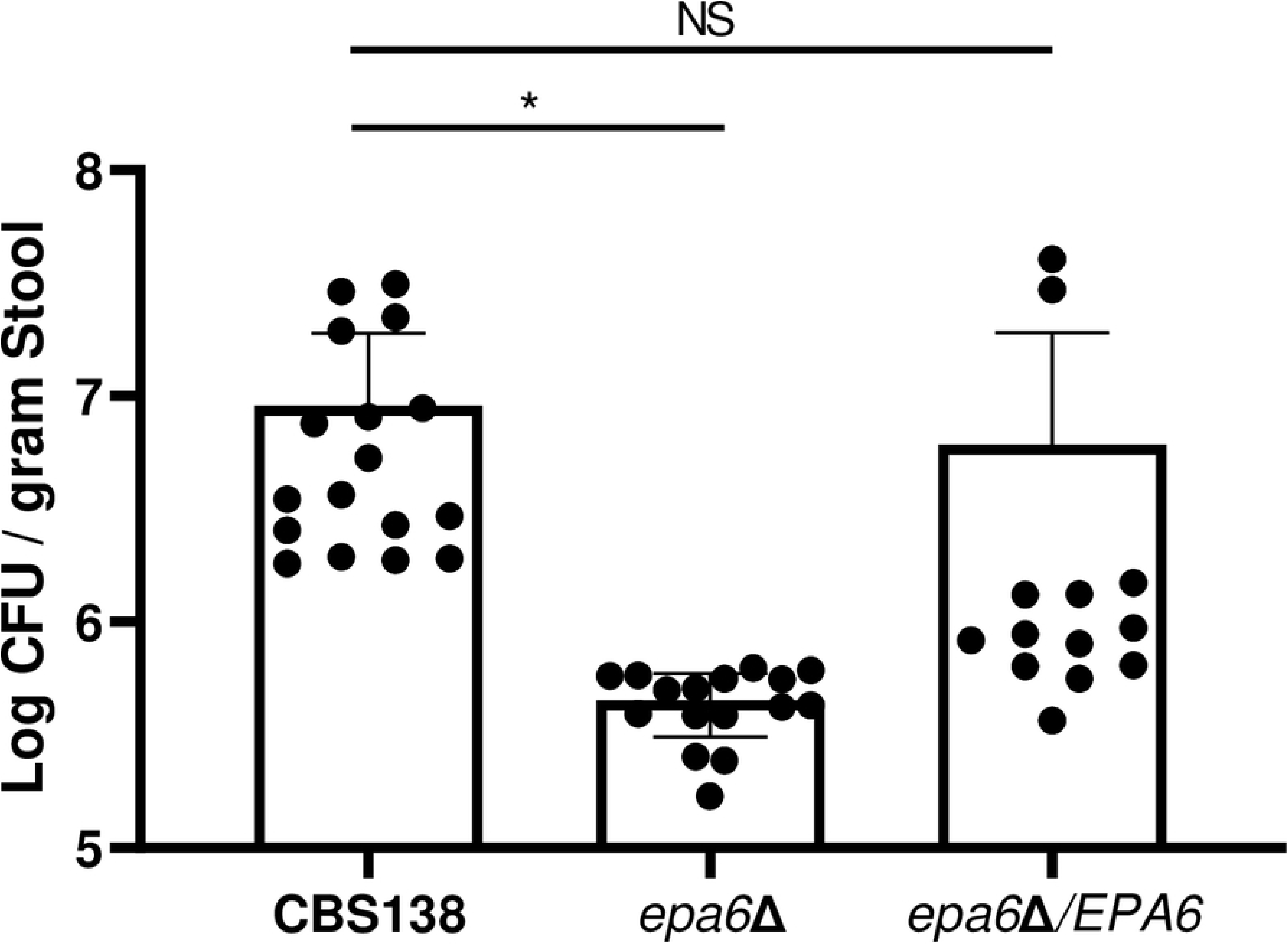
Stool fungal burden was significantly the lowest in mice infected with C. glabrata *epa6*Δ. The fungal burdens in stools collected 7 days after the inoculation of *C. glabrata* strains. Stool fungal burdens are indicated as the Log CFU/gram stool. All results are expressed as the mean ± standard error of the mean from five independent experiments with a total of 13–17 stool samples per group. * *p* < 0.05, NS, no significant difference.

**Fig 6.**
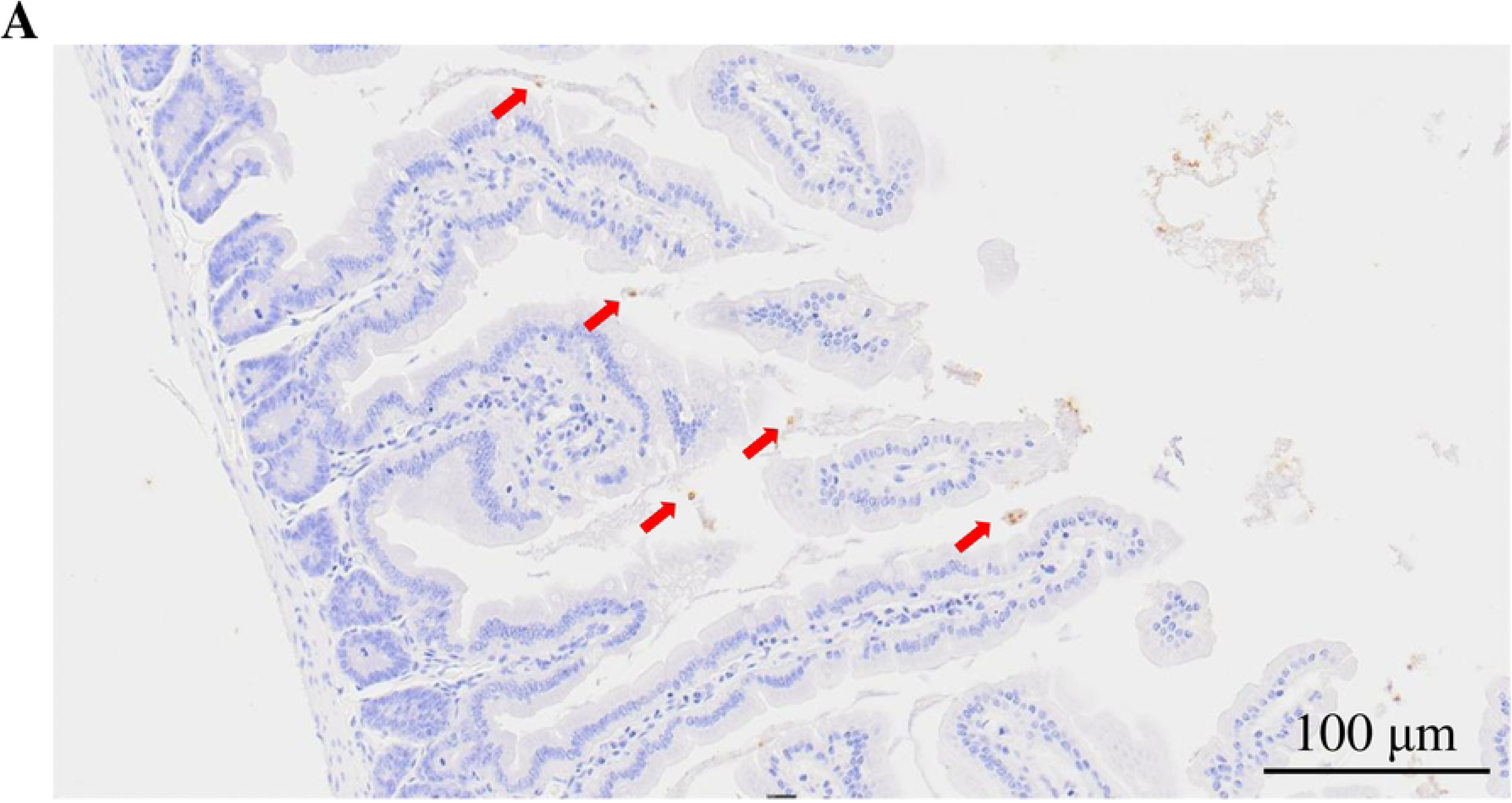
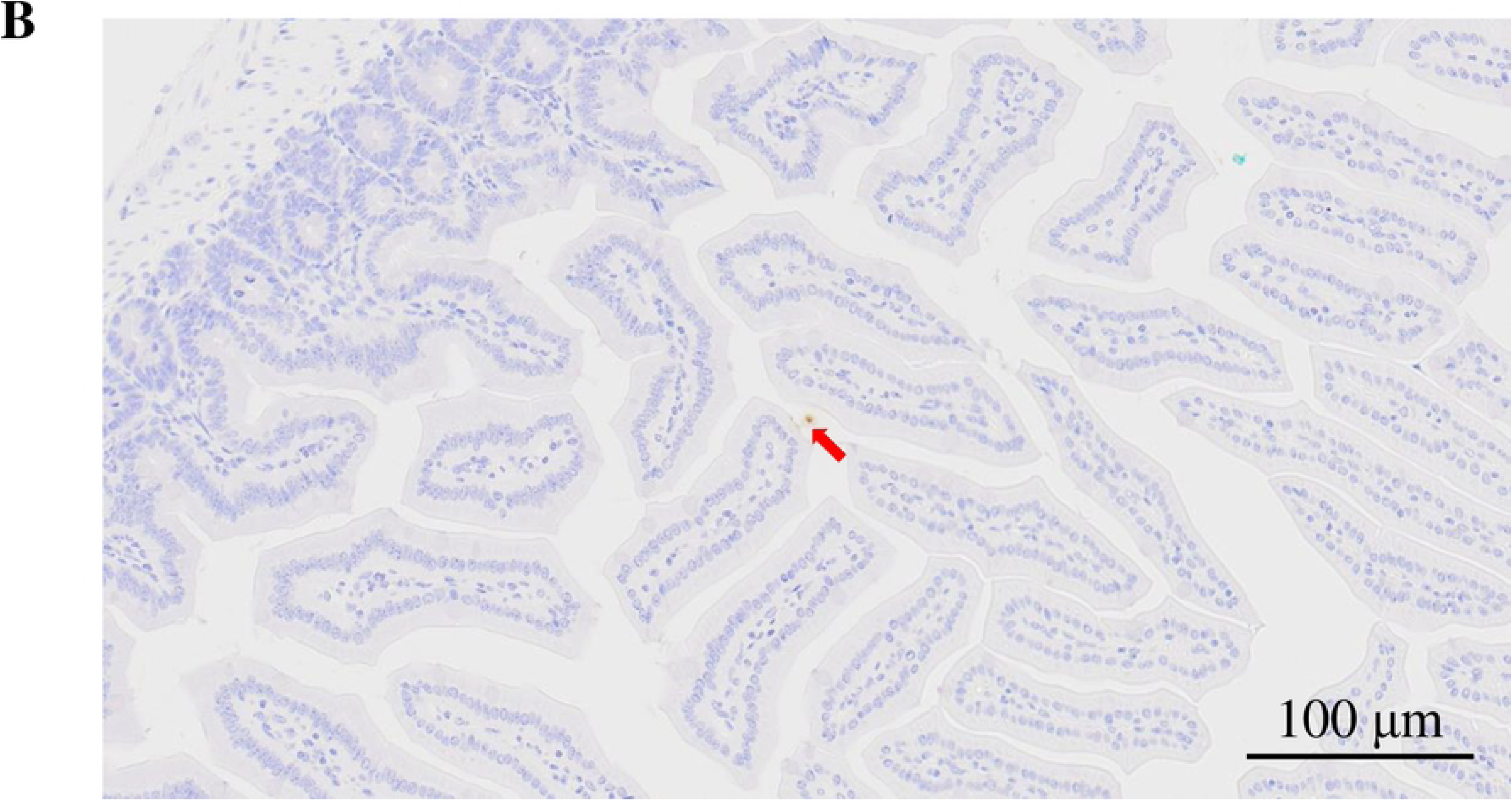
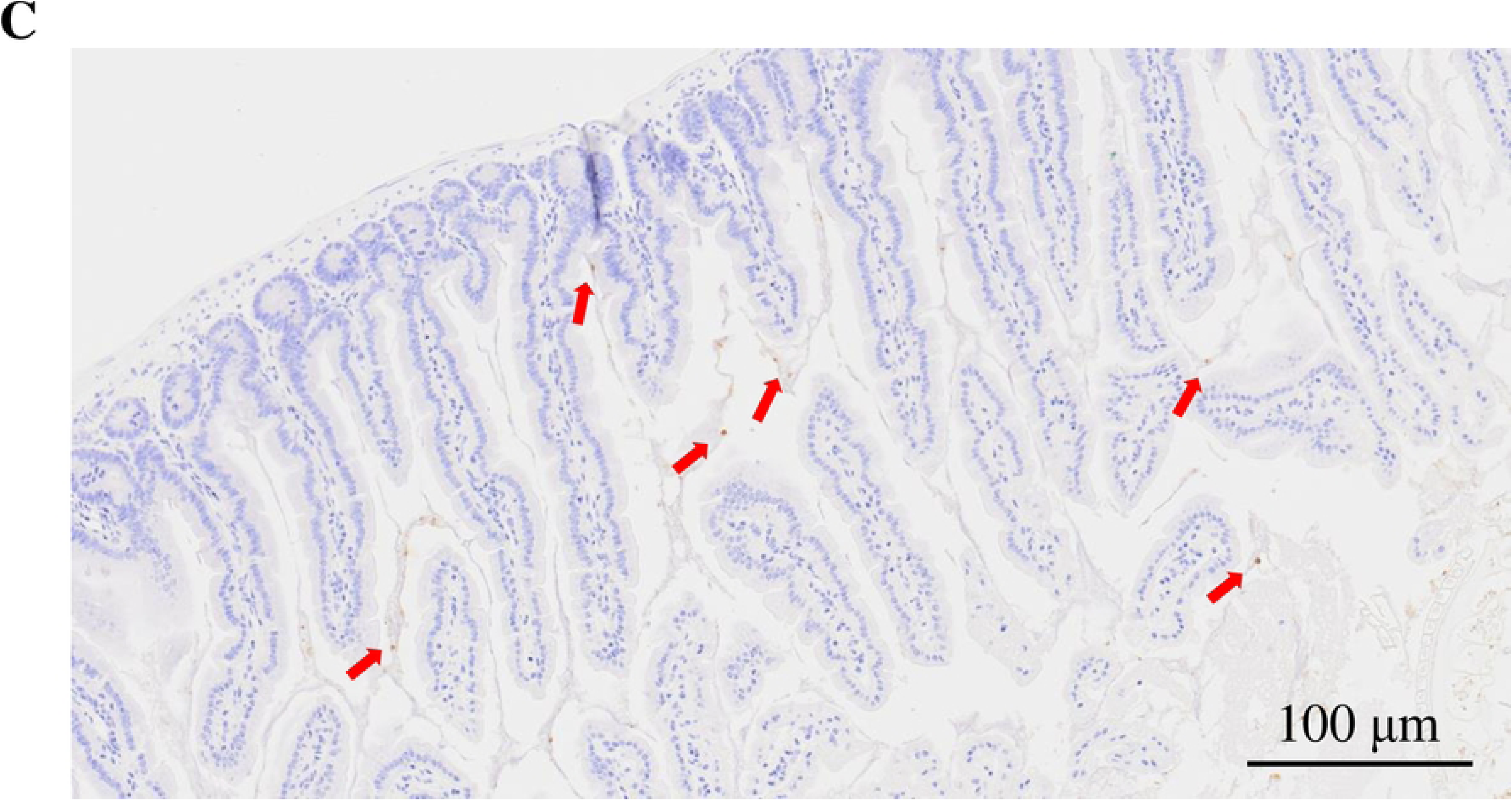
The *epa6*Δ mutant of *C. glabrata* colonized the murine colon less than CBS138 and *epa6*Δ/*EPA6*. (A–C) an anti-*Candida* antibody immunostaining of the intestine 7 days after inoculation with CBS138 (A), *epa6*Δ (B), and *epa6*Δ/*EPA6* (C). Arrows indicate *C. glabrata* yeast cells.

## Discussion

*Candida* species colonize the skin and mucocutaneous membranes where the environmental oxygen conditions differ. Therefore, oxygen levels surrounding *Candida* species could assumably affect their phenotypes. In this study, we showed that the adhesion capability of *C. glabrata* significantly increased under hypoxic conditions compared to those under normoxic conditions (Fig 1), and our RNA-seq analysis suggested that several adhesion-associated genes were up-regulated under anoxic conditions (Fig 2). Based on these results, we generated mutants for each of these adhesion genes and identified *EPA6* as the most critical gene in adhesion to Caco-2 cells (Fig 3). In addition, murine model experiments also demonstrated that *EPA6* was important for intestinal colonization (Figs 5 and 6). Taken together, we elucidated that oxygen concentration strongly affected the adhesive capability of *C. glabrata* and *EPA6* was the critical factor associated with this adhesive property.

Although the phenotype of *Candida* species in hypoxic conditions has not yet been investigated thoroughly, several studies have reported oxygen level-dependent phenotypic changes. For example, *C. albicans* inhibits hyphal formation under anaerobic conditions as a result of *EFG1* expression induction. This phenotype might be advantageous for commensalism in the host body [13]. Experiments under anaerobic conditions have often been performed using the AnaeroPack system (Mitsui Gas Co., Japan) or similar apparatus [17], exhibiting a limited working space and difficulty in measuring real-time oxygen concentrations. However, in the case of the Concept 400 Workstation (Ruskinn Technology Ltd, Leeds, UK) we used in this study, oxygen/carbon-dioxide concentrations and system temperature can be controlled in 0.1% and 0.1°C increments, respectively, by catalysts and gas mixtures, in addition to sufficient space for work. Using this instrument, experiments can also be conducted under microaerobic conditions to simulate in vivo environments. *C. tropicalis* reportedly promoted hyphal formation under microaerobic (5 vol% oxygen) compared to aerobic and anaerobic conditions with up-regulation of *ALS3* and *SAP1* [15]. These results indicate that each *Candida* species is likely to alter its phenotype depending on environmental oxygen levels. However, there are few reports describing the favorable oxygen conditions of *Candida* species. Our adhesion capability-related findings showed that hypoxic conditions were favorable for *C. glabrata* colonization and growth. In addition, as a novel insight, to our best knowledge, we elucidated that the anaerobic *C. glabrata* adhesion to intestinal cells was associated with the adhesive protein *EPA6*.

*EPA6* belongs to the *EPA* family [18, 19] of adhesion genes with expression levels changing drastically in response to the environment, normally negatively regulated by the Sir complex, yet de-repressed by specific external changes [20–22]. Further studies show that *EPA6* expression is positively regulated by Yak1p kinase acting through a subtelomeric silencing pathway and by the chromatin remodeling Swi/Snf complex, whereas it is negatively regulated by the transcription factor Cst6p [23]. Interestingly, our RNA-seq analysis revealed that the *SIR3* expression level was repressed upon 2 hours of anaerobic adhesion while remaining unchanged in aerobic adhesion (S3 Table), potentially suppressing the Sir complex followed by increased *EPA6* expression. Nicotinic acid deficiency and weakly acidic compounds are reportedly external stimuli that induce *EPA6* expression [24, 25]. Certain studies describe the relationship between *EPAs* and external stimuli [26]. However, none of these studies focused on the environmental oxygen conditions. Herein, we provided data indicating that hypoxic conditions were also related to *EPA6* expression and subsequent adhesive capabilities, presenting a novel aspect of external stimuli-induced *EPA* expression.

This study suggests that oxygen levels could alter *Candida* species phenotypes. Considering that *Candida* infections and propagation often occur in less oxygenated areas [7], evaluating the phenotype of *Candida* spp. under hypoxic conditions and providing an insight into host-pathogen interactions would be useful in various fields. This study indicates that Epa6p is the most important factor, especially under anaerobic conditions, for *C. glabrata* adhesion to the intestinal tract. Indeed, adhesion to host tissues or indwelling medical devices is an important first step in establishing fungal infection and is thus a crucial virulence factor [27]. If Epa6p function could be inhibited, it could contribute to reducing endogenous infection and biofilm formation. *C. glabrata* is known to be low-susceptible to azole antifungals. A novel antifungal agent type, such as one targeting these adhesive molecules, should thus be considered and further related studies would be necessary.

We met certain limitations in our experiments. First, our in vitro assays were simplistic and did not completely reflect actual human internal environments, such as interactions with bacteria, other fungi, and the host immune system. In our model, Epa6p was crucial for the adhesion to the intestinal tract under anaerobic conditions. However, other molecules might also be potentially involved in this process. Second, the oxygen level-sensing and eventual *EPA6* expression level-inducing mechanism of *C. glabrata* remain to be clarified. Mammals and plants have direct oxygen sensing mechanisms, such as the hypoxia-inducible factor-1 (HIF-1) pathway and group VII ethylene response factors regulated by the N-end rule pathway, respectively [28, 29]. Nevertheless, fungi are thought to use indirect sensing mechanisms, such as sterol homeostasis, mitochondrial reactive oxygen species, and nitric oxide productions [30]. Fungal sterol biosynthesis is so highly oxygen-consumptive that hypoxic conditions affect growth [31]. One hypothesis for increased adhesion under hypoxic conditions is that *C. glabrata* might deprive host cells of essential molecules that require oxygen for their biosynthesis. *C. glabrata* reportedly replaces or supplements its ergosterol with cholesterol recruited from host cells [32]. This property might be driven for survival under anaerobic conditions. Third, although invasive *Candida* infections are often a lethal problem in immunosuppressed patients [2], we did not use an immunosuppressed murine model. Analysis reflecting actual situations would be useful for elucidating pathogenesis, and further experiments have been planned.

In summary, to the best of our knowledge, this is the first study to describe anaerobic *C. glabrata* adhesion to intestine-derived Caco-2 cells. Our results suggest that anaerobic conditions promote *C. glabrata* adhesion and *EPA6* plays a significant role in hypoxic adhesion. If Epa6p function could be inhibited, it may contribute to reducing the colonization in the gut followed by endogenous infection. Evaluating the phenotype of *Candida* spp. under hypoxic conditions and providing an insight into host-pathogen interactions would be useful.

## Materials and methods

### Strains and media

*Escherichia coli* DH5α [F^−^ φ80*lac*ZΔM15 Δ(*lac*ZYA-*arg*F)U169 *rec*A1 *end*A1 *hsd*R17(r_K_ ^−^, m_K_ ^+^) *pho*A *sup*E44 λ^−^*thi*-1 *gyr*A96 *rel*A1] was used for plasmid propagation. Bacterial strains were grown in Luria-Bertani medium with ampicillin. The *C. glabrata* strains used in this study are listed in Table 1. The conditional YKU80 knockout strains and KUE200 were used to generate deletants [33]. All yeasts strains were usually grown in Yeast extract-Peptone-Dextrose (YPD) [1% Bacto Yeast Extract (Difco), 2% Bacto Peptone (Difco), and 2% glucose; pH 6.8] or Complete Supplement Mixture (CSM) [0.67% Bacto yeast nitrogen base without amino acids (Difco), 0.079% CSM (MP Biomedicals, Inc., OH, USA), 2% glucose, 40 mg/L additional adenine, and 60 mg/L additional histidine; pH 5.8] media, or 0.077% CSM without histidine. The solid medium was supplemented with 2% agar (Nacalai Tesque, Kyoto, Japan). Doxycycline for a Tet-off strain [16] was purchased from Sigma-Aldrich Chemical Co. (St Louis, MO).

**Table 1.** Candida glabrata strains.

| Strain | Origin |  | Reference |
| --- | --- | --- | --- |
| CBS138 | Reference strain |  | [34] |
| NIIDF0035502 | Clinical isolate from bloodstream infection |  | - |
| NIIDF0035508 | Clinical isolate from bloodstream infection |  | - |
| NIIDF0049901 | Clinical isolate from bloodstream infection |  | - |
| NIIDF0049902 | Clinical isolate from bloodstream infection |  | - |

|  | Parent | Genotype |  |
| --- | --- | --- | --- |
| 2001H | 2001U | $\Delta his3::ScURA3 \Delta ura3$ | [35] |
| 2001HT | 2001TU | $\Delta trp1 \Delta his3::ScURA3 \Delta ura3$ | [35] |
| KUE100 | 2001H | $\Delta his3::ScURA3 \Delta ura3 yku80::SAT1 flipper$ | [33] |
| KUE200 | 2001HT | $\Delta trp1 \Delta his3::ScURA3 \Delta ura3 yku80::SAT1 flipper$ | [33] |
| *epa6*Δ | KUE200 | $\Delta trp1 \Delta his3::ScURA3 \Delta ura3 yku80::SAT1 flipper epa6::CgHIS3$ | this study |
| *epa6*Δ/*EPA6* | *epa6*Δ | $\Delta trp1 \Delta his3::ScURA3 \Delta ura3 yku80::SAT1 flipper CgHIS3::CgHIS3-EPA6-CgTRP1$ | this study |
| *epa2*Δ | KUE200 | $\Delta trp1 \Delta his3::ScURA3 \Delta ura3 yku80::SAT1 flipper epa2::CgHIS3$ | this study |
| *awp1*Δ | KUE200 | $\Delta trp1 \Delta his3::ScURA3 \Delta ura3 yku80::SAT1 flipper awp1::CgHIS3$ | this study |
| *awp2*Δ | KUE200 | $\Delta trp1 \Delta his3::ScURA3 \Delta ura3 yku80::SAT1 flipper awp2::CgHIS3$ | this study |
| *pwp3*Δ | KUE200 | $\Delta trp1 \Delta his3::ScURA3 \Delta ura3 yku80::SAT1 flipper pwp3::CgHIS3$ | this study |
| *cagl0j00253gΔ* | KUE200 | *Δtrp1 Δhis3::ScURA3 Δura3 yku80::SAT1 flipper cagl0j00253g::CgHIS3* | this study |
| ACG4 | 2001HT | *his3 trp1 PScADH1::tetR::GAL4AD::TRP1* | [36] |
| HET80 | ACG4 | *yku80 ILV2(P182L)* | [36] |
| HETS202 | HET80 | *his3 trp1 PScADH1::tetR::GAL4AD::TRP1 yku80::SAT1 flipper* | [36] |
| Tet-off CAGL0J02552g | HETS202 | same as ACG4 but *cagl0j02552g::HIS3-97t-CAGL0J02552g* | this study |

### Adhesion assays

Adhesion assays were performed using the Caco-2 colorectal carcinoma-derived cell line. Cells were grown in 12-well plastic dishes in Roswell Park Memorial Institute medium supplemented with 10% fetal bovine serum and antibiotics (penicillin/streptomycin). Cells were allowed to differentiate in monolayers for 7–10 days at 37°C with 5% CO_2_. *C. glabrata* strains were cultured overnight to stationary phase at 37°C in YPD broth and used to infect Caco-2 cells at a density of 1×10^5^ cells per well under aerobic (21 vol% oxygen), microaerobic (5 vol% oxygen), and anaerobic (0 vol% oxygen) conditions in a Concept 400 Workstation (Ruskinn Technology Ltd, Leeds, UK). After incubation for two hours at each oxygen level, nonadherent cells were removed by three washes in phosphate-buffered saline (PBS). Adherent cells were recovered by lysing the monolayer in 0.1% Triton X-100 and quantified by counting colony-forming units (CFUs) after 24 hours of incubation at 37°C on YPD plates. The percentage of adherent cells, referred to as %adherence, was calculated as CFUs recovered by lysis of the monolayer divided by the CFUs added.

### Preparation of RNA sequencing (RNA-seq) libraries, sequencing, and data analysis

For total RNA extraction, *C. glabrata* (CBS138) was grown overnight in the stationary phase under aerobic or anaerobic conditions and recovered two hours after adhesion to Caco-2 cells under each oxygen level. RNA was then purified and resuspended with SMART-Seq v4 Ultra Low Input RNA Kit (Takara Bio Inc., Shiga, Japan) according to the manufacturer’s instructions for mRNA amplification using 5’ template switching polymerase chain reaction (PCR). Amplified cDNA was fragmented and appended with dual-indexed barcodes using Illumina Nextera XT DNA Library Prep kits (Illumina, San Diego, CA). Libraries were validated on an Agilent 4200 TapeStation (Agilent Technologies, Santa Clara, CA), pooled, and sequenced on an Illumina NovaSeq 6000 platform (Illumina, San Diego, CA). Read sequences obtained by sequence analysis were mapped to a reference genome sequence of *C. glabrata* CBS138 from the GENCODE database version 35. Transcript abundance was estimated as transcripts per million (TPM) based on the length of the gene and the number of reads mapped to the gene.

### Adhesin mutant construction

The knockout cassette used to replace *EPA6*, *EPA2,* or *AWP2* ORF with *HIS3* was amplified from the plasmid pHIS916 [33] using the primer pairs EPA6DF/EPA6DR, EPA2DF/EPA2DR, or AWP2DF/AWP2DR, respectively. As for *PWP3*, three fragments were joined via overlap PCR using the primer pair PWP3DF/PWP3DR, consisting of two fragments (*PWP3* 5’ and 3’ flanking regions) amplified from CBS138 genomic DNA using the primer pairs PWP3-5F/PWP3-5R and PWP3-3F/PWP3-3R, as well as a fragment *HIS3* amplified from pHIS906 using the primer pair PWP3HDF/PWP3HDR.

To construct an *AWP1* or *CAGL0J00253g* knockout cassette, the In-Fusion cloning technology was performed as described previously [37]. Briefly, the *HIS3* cassette was amplified from pHIS906 using the primer pair HIS3F/HIS3R. Five hundred-base pair (bp) *AWP1* 5’ and 3’ flanking regions were amplified from CBS138 genomic DNA with the primer pairs AWP1-5F/AWP1-5R and AWP1-3F/AWP1-3R, respectively. pHIS906 was linearly PCR-amplified using the primer pair 906LF/906LR, followed by digestion with DpnI. To introduce compatible ends for cloning, the primer pair AWP1-5F/AWP1-3R was used containing 15-bp-long 5’ overhangs identical to the linearized pHIS906. Similarly, the primer pair AWP1-5R/AWP1-3F contained 15-bp 5’ overhangs identical to HIS3F/HIS3R sequences. Next, all the three amplicons (the 5’ flanking region, the *HIS3* cassette, and the 3’ flanking region) and the linearized pHIS906 were fused using the In-Fusion enzyme (TAKARA, Shiga, Japan) to create the pHIS3-AWP1 plasmid. A similar method was applied to construct the pHIS906-00253g plasmid using a 500-bp *CAGL0J00253g* 5’ flanking region and a 500-bp 3’ flanking region, amplified from CBS138 genomic DNA with the primer pairs 00253-5F/00253-5R and 00253-3F/00253-3R, respectively. Using these plasmids as templates, the knockout cassette to replace the *AWP1* or *CAGL0J00253g* ORF with *HIS3* was amplified from the pHIS3-AWP1 or pHIS906-00253g plasmids by the primer pairs AWP1DF/AWP1DR or 00253DF/00253DR, respectively.

Each amplified knockout cassette was used for the transformation of KUE200 [33]. The disruption of these genes was PCR-verified using the primer pairs pTET12F/EPA6CHR, pTET12F/EPA2CHR, pTET12F/AWP2CHR, pTET12F/PWP3CHR, pTET12F/AWP1CHR, or pTET12F/00253CHR for *EPA6, EPA2, AWP2*, *PWP3*, *AWP1,* or *CAGL0J00253g*, respectively.

To generate a *CAGL0J02552g* Tet-off strain, *C. glabrata* was transformed as previously described [33]. A DNA cassette was PCR-amplified from the pTK916-97t plasmid [38] using the primer pair 02552tetF/02552tetR. The resulting product was transformed into HETS202 cells [38]. To confirm the integration of the cassette, it was PCR-verified using the primer pair 02552tetCHF/02552tetCHR. S1 Table summarizes the sequences of all primers used in this study.

### Wild-type allele reintegration into *EPA6* deletants

The DNA fragment harboring the *EPA6* ORF from a 1000-bp *EPA6* 5’ flanking region to a 500-bp 3’ flanking region was amplified from the CBS138 genome using the primer pair EPA6cF/EPA6cR. The amplified *EPA6*-containing fragment was cloned into pTi-comp plasmid [35] by In-Fusion cloning as described above. Briefly, the pTi-comp plasmid was PCR-amplified linearly using the primer pair TICOMPLF/TICOMPLR, followed by digestion with DpnI. To introduce compatible ends for cloning, we used the primer pair EPA6cF/EPA6cR containing 15-bp-long 5’ overhangs identical to the linearized pTi-comp. Next, the *EPA6* cassette and the linearized pTi-comp were fused using the In-Fusion enzyme (TAKARA, Shiga, Japan) to create the pTi-comp-EPA6 plasmid. Using this plasmid as a template, the insertion fragment was amplified with the primer pair EPA6REVF/EPA6REVR. The resulting DNA fragment was introduced into the *HIS3* gene-replaced locus on the deletant chromosome by end-in type recombination [39]. The accurate insertion of the amplified fragment into the correct chromosomal locus was PCR-confirmed using the primers EPA6RCHF/EPA6RCHR. S1 Table summarizes the sequences of all primers used in this study.

### Reverse transcription-quantitative real-time (RT-q) PCR-based gene expression analysis

The relative *EPA6* mRNA expression level was assessed by RT-qPCR at the time of inoculum to Caco-2 cells and two and eight hours after aerobic or anaerobic adhesion. The adherent *C. glabrata* was recovered for each time point using a cell scraper. Total RNA was extracted and purified from each sample using beads and ISOGEN (Nippon Gene, Toyama, Japan), RNeasy Mini Kit (Qiagen, Tokyo, Japan), and RNase-free DNase I (Qiagen, Tokyo, Japan) following the manufacturers’ instructions. The resulting RNA samples were processed for reverse transcription using the ReverTra Ace qPCR RT Master Mix kit (Toyobo, Osaka, Japan). RT-PCR was performed in an Mx3000P real-time PCR system (Agilent, Santa Clara, CA) using SYBR Premix ExTaq (Takara, Shiga, Japan). The relative *EPA6* expression level was normalized by comparison to the expression of 18S-rRNA. These experiments were performed in biological triplicates. S1 Table summarizes the sequences of all primers used in this study.

### Mice

Male, 6−7-week-old C57BL/6J mice were purchased from Japan SLC, Inc. (Shizuoka, Japan) and maintained under specific pathogen-free conditions at the National Institute of Infectious Diseases in Japan. All experiments were approved by the Animal Care and Use Committee of the National Institute of Infectious Diseases. All staff members, including the participants of this investigation, were sufficiently educated in animal care and handling before animial experiments. Protocols were designed for minimizing animal suffering and limiting the number of animals used in experiments (Approval numbers: 121129).

### Yeast strains and murine model colonization

*C. glabrata EPA6* deletant (*epa*6Δ), reintegrated (*epa*6Δ/*EPA6*), and reference (CBS138) strains were used in this study (Table 1). For the *C. glabrata* colonization murine model, mice were administered with an antibiotics cocktail (consisting of vancomycin, gentamicin, kanamycin, and metronidazole at 45, 35, 400, and 215 mg/L, respectively, as well as colistin at 850 U/L, as described previously [40]) in their drinking water for more than a week before inoculation with *C. glabrata* [40]. All *C. glabrata* strains were grown at 37°C on YPD agar, then in YPD broth for one day and 12−18 hours, respectively. The yeast cells were collected, washed, and resuspended in sterile PBS at a concentration of approximately 5.0 × 10^8^−1.0 × 10^9^ CFU/mL. Mice were intragastrically inoculated with 100 μL (approximately 5.0 × 10^7^−1.0 × 10^8^ CFU/mouse) via a sterile feeding needle. Seven days after inoculation, fresh stools from each mouse were collected for evaluating gastrointestinal colonization. Stools were homogenized in sterile PBS, then the homogenates were serially diluted and plated on YPD agar with penicillin/streptomycin to inhibit bacterial growth. The fungal burden was determined by counting CFUs after 24 hours of incubation at 37°C.

### Infected mouse gut histopathological analysis

For histopathological analysis, guts were removed 7 days after *C. glabrata* inoculation, fixed in 10% neutral-buffered formalin, dehydrated with ethanol, and embedded in paraffin following standard procedures. Four-μm tissue sections were mounted onto glass slides (Matsunami Glass, Osaka, Japan) and stained with hematoxylin and eosin, Grocott methenamine silver, or an anti-*Candida* antibody (in-house rabbit polyclonal antibody against *C. albicans*). Histological examinations were performed using light microscopy.

### Statistical analysis

Statistical analyses were performed using GraphPad Prism, version 8 (GraphPad Software, La Jolla, CA). One-way analysis of variance (ANOVA) with a Tukey multiple comparison posttest was used to analyze more than 2 groups. P-values of less than *p <* 0.05 were considered statistically significant for all tests.

## Acknowledgments

The authors would like to thank Ms. Yuko Sato for her technical assistance with pathological analysis.

## Author contributions

Conceptualization: Takayuki Shinohara and Yoshitsugu Miyazaki.

Data curation: Takayuki Shinohara.

Formal analysis: Takayuki Shinohara.

Funding acquisition: Masahiro Abe, Harutaka Katano, and Yoshitsugu Miyazaki.

Investigation: Takayuki Shinohara and Harutaka Katano.

Methodology: Takayuki Shinohara, Harutaka Katano, and Hiroji Chibana.

Project administration: Yoshitsugu Miyazaki.

Resources: Yoshitsugu Miyazaki.

Supervision: Yoshitsugu Miyazaki.

Revision and editing: Masahiro Abe, Sota Sadamoto, Minoru Nagi, Harutaka Katano, Hiroji Chibana, and Yoshitsugu Miyazaki.

## Meeting(s) where the information has previously been presented

We presented this study at the 97th Annual Meeting of the Japanese Association for Infectious Diseases.

## Supporting information

**S1 Table.** Primers used in this study.

**S2 Table.** RNA-seq analysis. The TPM values of *C. glabrata* extracted at the time of inoculum and two hours after Caco-2 cell adhesion under anaerobic and aerobic conditions, respectively.

**S3 Table.** The TPM value fold changes of adhesion genes upon two hours under aerobic and anaerobic adhesion.

